# Molecular dynamics simulations of hydrophobic peptides that form β-hairpin structures in solution

**DOI:** 10.1101/2021.10.08.463620

**Authors:** Tushar Ranjan Moharana, Ramakrishnan Nagaraj

**Author notes:** Author for correspondence: R. Nagaraj, CSIR-Centre for Cellular and Molecular Biology, Uppal Road, Hyderabad 500 007, India.

## Abstract

Peptides designed with residues that have high propensity to occur in β-turns, form β-hairpin structures in apolar solvents as well in polar organic solvents such as dimethyl sulfoxide (DMSO), methanol and varying percentages of DMSO in chloroform (CHCl_3_). Presumably due to limited solubility, their conformations have not been investigated by experimental methods in water. We have examined the conformations of such designed peptides that fold into well-defined β-hairpin structures facilitated by β-turns, in the crystalline state and in solution, by Molecular Dynamics Simulations (MDS). The peptides fold into β-hairpin structures in water, starting from extended conformation. In DMSO, folding into β-hairpin structures was not observed, starting from extended conformation. However, when the starting structure is in β-hairpin conformation, unfolding is not observed during MDS in DMSO. Water clearly favours folding of short, hydrophobic peptides into β-turn and β-hairpin conformations from extended structures. DMSO does not have a denaturing effect on short, hydrophobic peptides.

## Introduction

Short linear peptides do have the ability to adopt folded conformation in solution, although it has been generally believed that short peptides are unstructured in solution [1-10]. Nascent ordered structures have been proposed for peptides in aqueous solution, based on Nuclear Magnetic Resonance (NMR) data [10]. Distinct β-turn and β-hairpin conformations have been proposed for short designer peptides, based on detailed analysis of their two dimensional (2D) NMR spectra [1-9]. These peptides adopt β-hairpin conformation with a well-defined β-turn in apolar solvents such as chloroform (CHCl_3_). Few of them fold into β-turn and β-hairpin conformation in hydrogen bonding solvents such as dimethyl sulfoxide (DMSO), methanol and water. These peptides are composed of hydrophobic and Phe (F) residues with tight β-turns formed by pX where p is D-Pro and X is Gly (G), Ala (A), or α-aminoisobutyric acid (Aib). A triple stranded structure has also been observed in a model peptide composed of entirely of hydrophobic residues and F with two β-turns formed by pG [3]. Of particular interest in the observation of distinct β-turn and β-hairpin structure for a model peptide t-butyloxycabonyl (Boc)-Leu-Val-Val-D-Pro-Gly-Leu-Val-Val-OMe, where OMe is methylester, in DMSO [1], as observed in the crystalline state [11]. This solvent is generally used to dissolve aggregating peptides such as amyloidogenic peptides to break up any secondary structures [12]. Infact, several peptides that are not composed of hydrophobic amino acids do not adopt folded structures in DMSO [13-16].

In an effort to set insights into the conformational dynamics of model peptides that exhibit distinct conformations in solution and crystalline state [2, 3, 11, 17, 18], we have carried out Molecular Dynamics Simulations (MDS) of peptides whose structures have been characterised in detail in solution and in the crystalline state. The peptides studied (Note: one letter code has been used as follows: Leu, L; Val. V; D-Pro, p; Gly. G; Phe, F) were Acetyl(Ac)-LVVpGLVV-NH_2_-**1**; Ac-LFVpGLFV-NH_2_-**2**; Ac-LFVpGLVLApGFVV-NH_2_-**3**; Ac-LFVpALFV-NH_2_-**4**. The C-terminal was amidated and is indicated as -NH_2_. All the peptides have the β-turn promoting sequence pG and pA. Peptide **3** is characterized by two turns [3]. While the peptides fold into β-turn and β-hairpin structure in water, they show considerable structural flexibility in DMSO.

## Methods

### Initial structure preparation

Atomic coordinates of peptides atoms without hydrogens, in extended conformation, was prepared by Discovery Studio 2019. Dihedral angles of all the amino acids were kept at 180° except D-Pro (Ф=60°, Ψ=180°). Acetyl and amide groups were attached to N and C terminus of each peptide respectively. Atom names were modified manually to be consistent with the forcefield convention. Hydrogen atoms were added by pdb2gmx module of GROMACS. Protonation states were modelled as expected at pH 7.

### System preparation

Peptide was kept at the centre of a cubic box. Dimension of the box was such that there is at least 1 nm distance between any atom of the peptide and surface of the box during folding simulation and 1.5 nm during unfolding simulation. The box containing peptide was solvated by solvent box, pre-equilibrated at 25°C and one bar. Finally, solvent molecules were allowed to diffuse around the solute atoms. Position restrain (1000kJ mol^-1^nm^-2^) was applied on heavy atoms of the peptide, during this process, to avoid any structural deformation.

For MDS in aqueous medium, pre-equilibrated water box containing 216 water molecules was used as solvent. For MDS in DMSO, pre-equilibrated DMSO box containing 250 molecules of DMSO was used as solvent.

### D-Pro and DMSO topology

Parameters of D-Pro were derived from that of proline by altering the dihedral angle Ф. DMSO parameters were from Vishnyakov et al. [19]. Atom types of sulphur, oxygen and carbons were assigned as SDMSO, ODMSO and CDMSO respectively. Methyl groups of DMSO were modelled by united atom approach. Partial atomic charges, bonded and non-bonded interactions were taken from Vishnyakov et al. [19].

### Molecular dynamics simulation

MDS was carried out using GROMACS package (version 5.0.4) [20]. GROMOS54a7 united atom forcefield [21] was used along with simple point charge water model [22] as recommended. System was equilibrated by 50000 steps of steepest gradient, 100 ps of nVT and 100 ps of nPT prior to actual MDS. Simulations were carried out for 100 ns in nPT ensemble. All the simulations were carried out at standard temperature (25°C) and pressure (1 bar). During MDS, pressure was maintained by Parrinello-Rahman barostat [23] and temperature was maintained by velocity-rescaling thermostat [24] with time constants of 2 ps and 0.1 ps respectively. The Lennard-Jones interactions and short range electrostatic interaction were truncated at 1.2 nm. Long term electrostatic interactions were handled by Particle Mesh Ewald (PME) method [25] with PME order 4 and fourier grid spacing at 0.16 nm. SETTLE algorithm was used to maintain water geometry [26]. All other bonds were constrained through LINCS algorithm [27] with lincs order 4. Edge of the simulation box was treated with periodic boundary condition. Calculation was done for a time step of 2 fs. Neighbour list was updated after every 10th step. Coordinates were saved in an interval of 10 ps.

### Visualization and analysis

Traces of distances and hydrogen bond count were generated from the trajectory using gmx-distance and gmx-hbond modules of GROMACS respectively. RMSD were calculated from the starting structure using gmx-rms module. Structures were generated using Discovery Studio 19 v19.1.0.18287. The dihedral angle plots were generated using VMD software.

## Results and Discussion

MDS were carried out on Ac-LVVpGLVV-NH_2_-**1**, Ac-LFVpGLFV-NH_2_-**2**, Ac-LFVpGLVLApGFVV-NH_2_-**3** and Ac-LFVpALFV-NH_2_-**4**. The structures of **1, 2** and **4** with t-Butyloxycarbonyl at the N-terminus and methyl ester at the C-terminus, was determined by X-ray crystallography (Figure 1).

**Figure 1:**
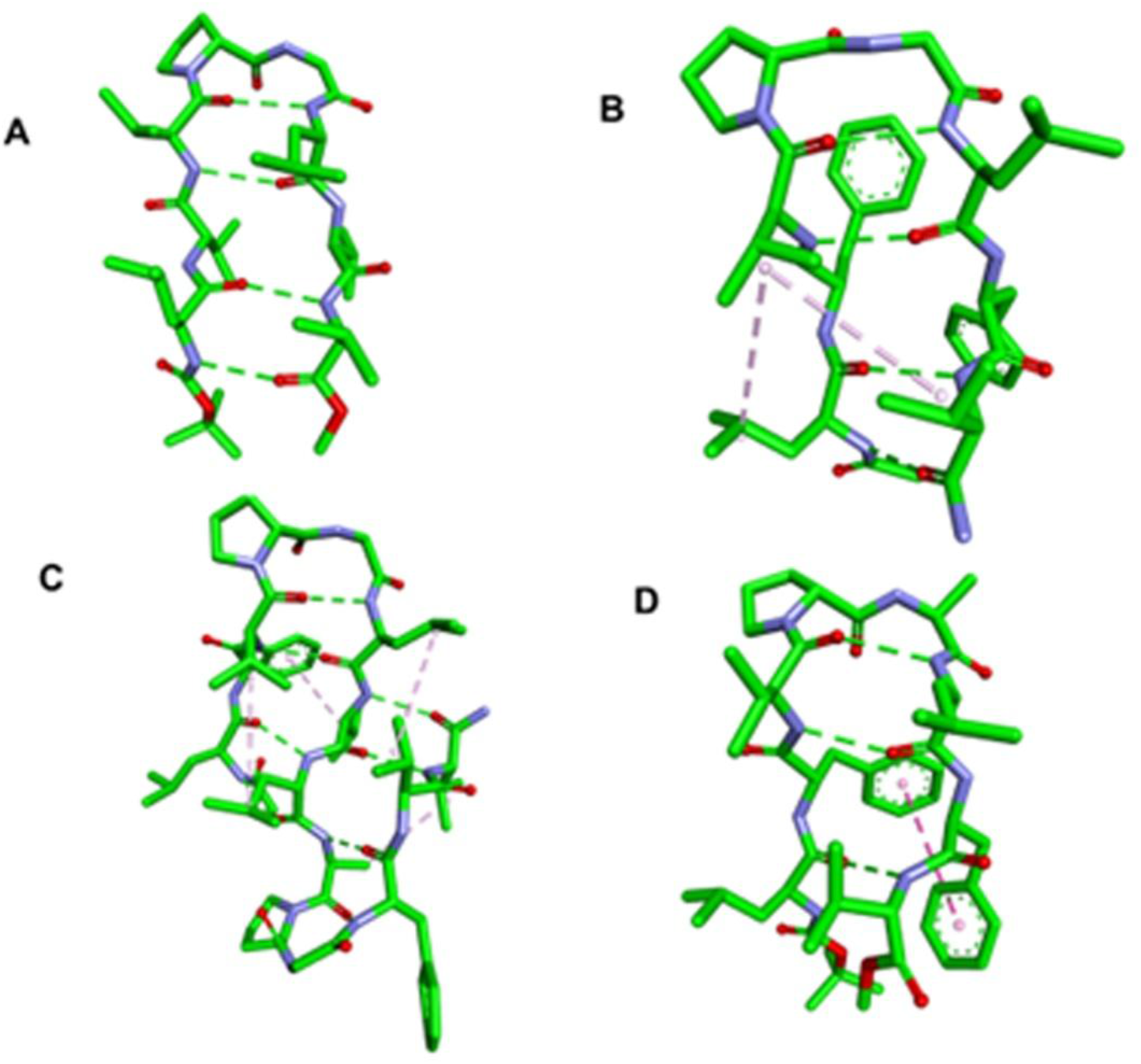
Structures of peptides. (A) Boc-LVVpGLVV-OMe [11]; (B) Boc-LFVpGLFV-OMe [17]; (C) Ac-LFVpGLVLApGFVV-NH2 modelled from the structure Boc-LFVpGLVLApGFVV-OMe [3] and (D) Boc-LFVpALFV-OMe [18]. Backbone hydrogen bonds, interaction between aromatic rings and aliphatic interactions are indicated by dashes. D-amino acid is indicated by p.

Peptide **3** was modelled based on the structures of **1, 2** and **4**. The β-turns are Type 2′ (T_2_′) in all the peptides. MDS was carried out from fully extended starting structures with Ф, Ψ=180° except for p where Ф was fixed 60°. MDS was carried out in water and DMSO. The distributions of Ф, Ψ during MDS in water for p and G in the peptides are shown in Figure 2.

**Figure 2:**
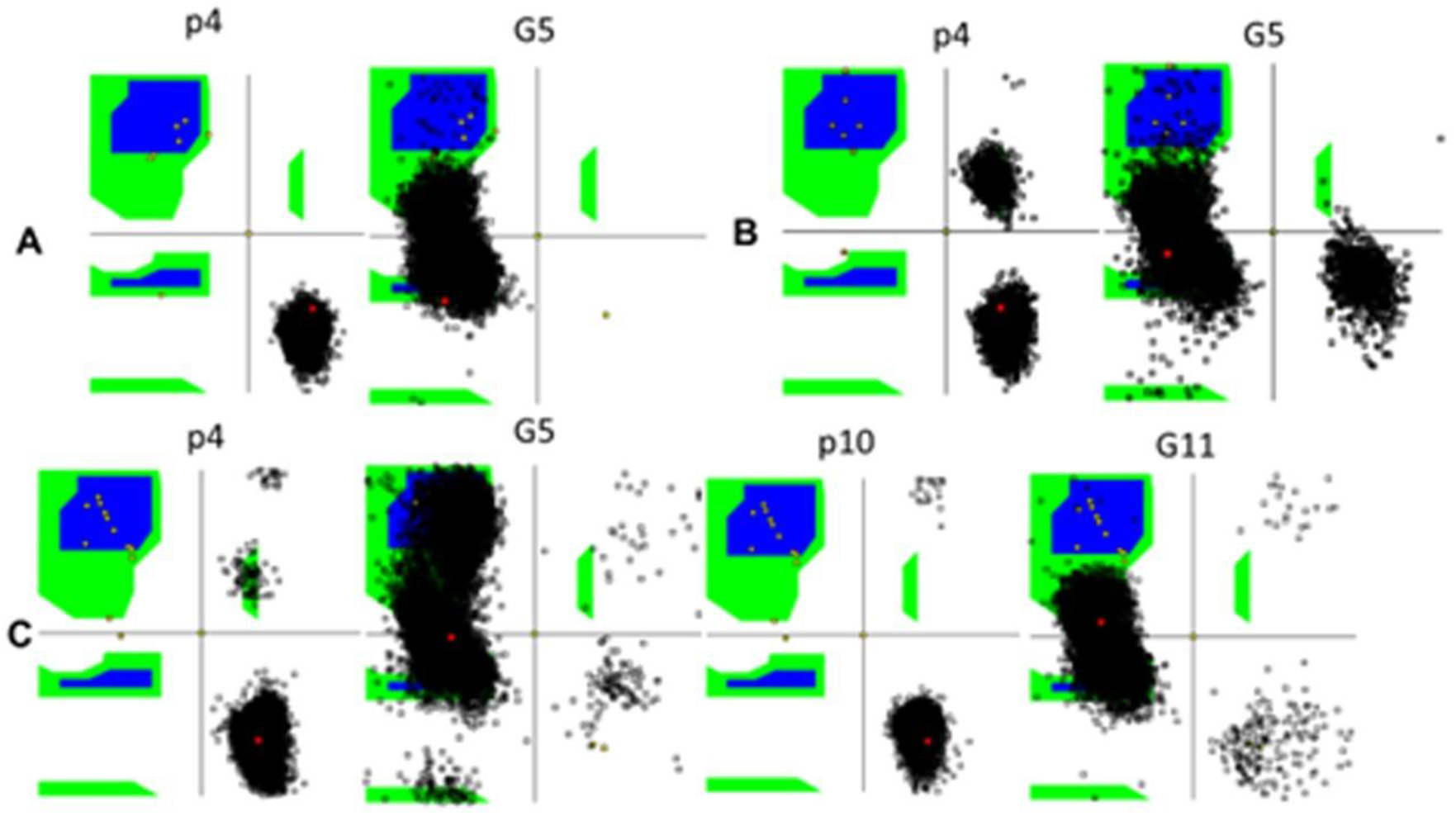
Distribution of Ф, Ψ for p and G in peptides. (A) **1**, (B) **2** and (C) **3** during the course of MDS. p4, G5 are D-Pro and Gly at positions 4, 5 in peptides **1-3**. p10, G11 are D-Pro and Gly at positions 10,11 in **3**. Black dots indicate Φ, Ψ values for the indicated amino acids during MDS.

The Ф, Ψ distribution for p is in the region ofT_2_′ β-turn for **1, 2** and **3**. In **2**, a small population occurs in Type 1′ (T1′) β-turn conformation (p with positive Ф, Ψ values). Ф, Ψ values of G show a broader spread. However, the area corresponding to T_2_′ β-turn is populated significantly. In **3**, the dihedral angles of G in the first turn formed by pG sample a larger area as compared to G in the second pG turn, indicating that it is more flexible as compared to the second turn.

The backbone RMSD, distances between N and C-terminal ends of the peptides and β-turns are shown in Figure 3. They indicate rapid folding of the peptides. The distances between ends of the four residues that form the β-turn indicate there are fluctuations in the β-turn geometry which is also reflected in the Ф, Ψ values of the p, G residues in the peptides, particularly for G.

**Figure 3:**
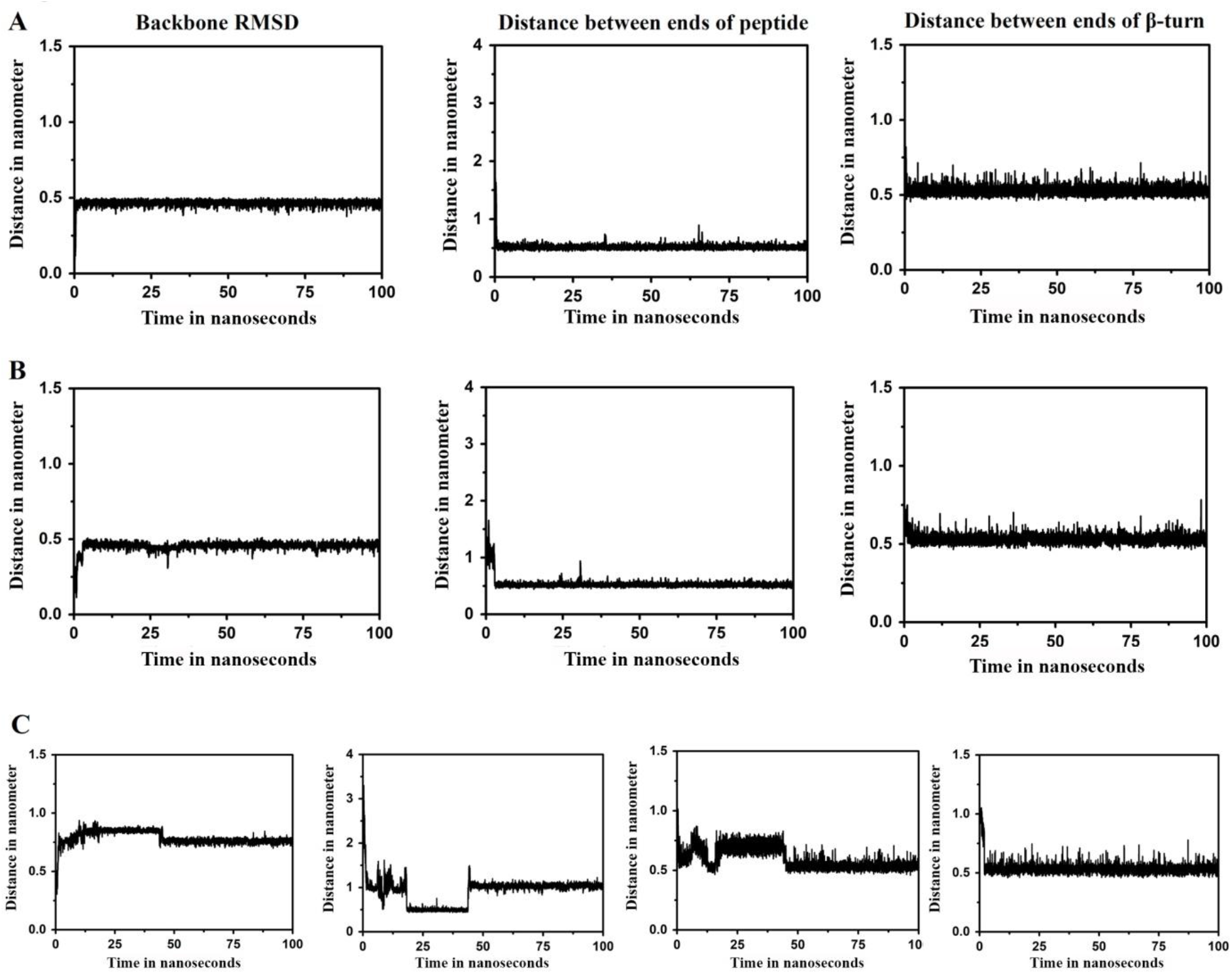
Backbone RMSD, distance between Cα of N and C-terminal amino acids and distance between Cα_I_ and Cα_i+4_ of residues forming β-turns in peptides. (A) **1**, (B), **2** (C) **3**.

Snapshots of structures during the course of MDS are shown in Figure 4. Peptide **1** folds early at 10 ns. Peptides **2** and **3**, fold into β-hairpin structures at 50 ns. The structures are stabilized by backbone hydrogen bonds and hydrophobic interactions. The hydrogen bonds are formed early during MDS. The conformations of **1** and **3** have been examined in CHCl_3_ [1, 3]. Conformations as shown in Figure 1 have been proposed in this apolar solvent. MDS clearly indicate that the peptides **1-3** can fold into β-hairpin conformation in water.

**Figure 4:**
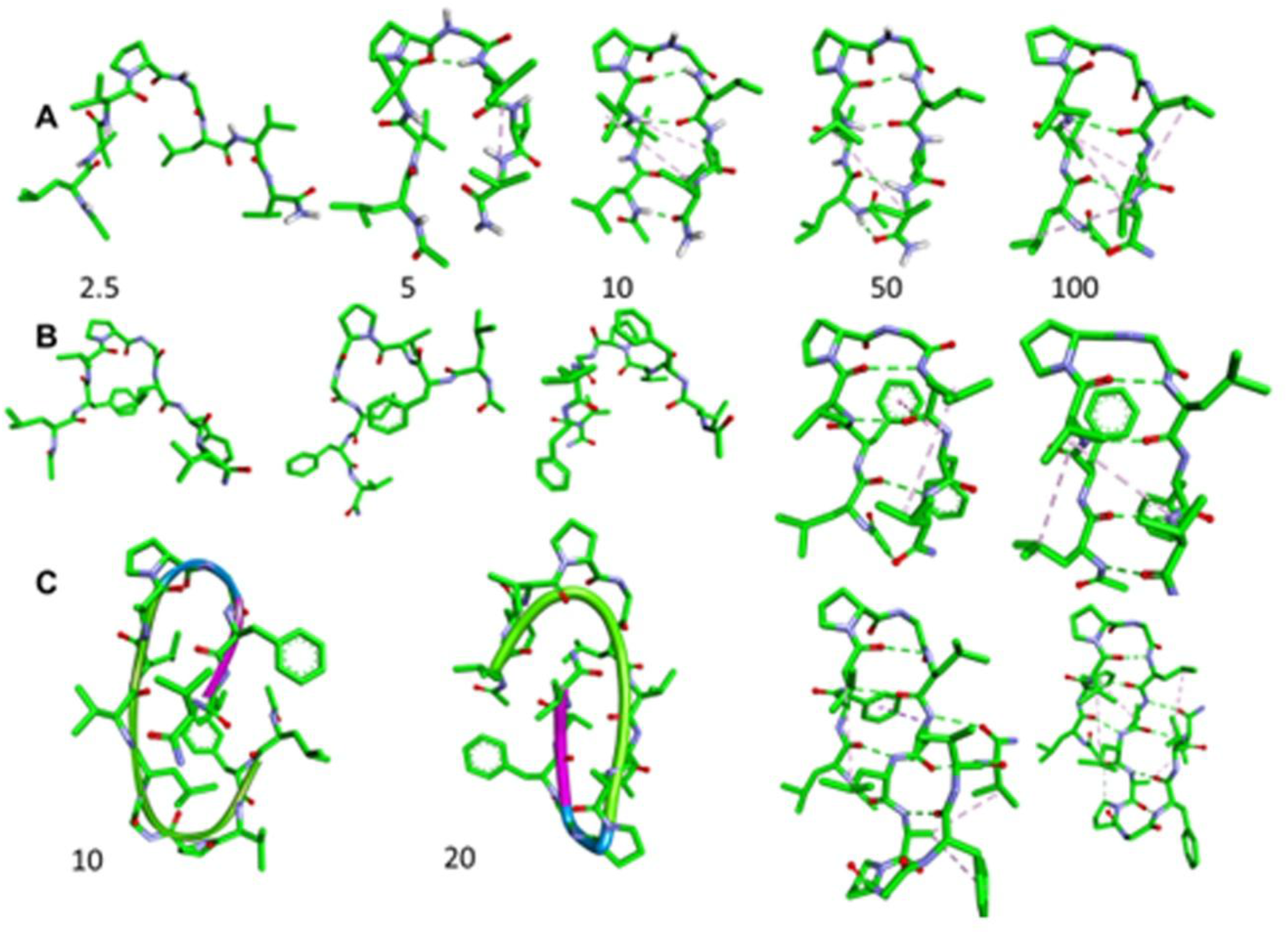
Snap shots of peptides at different times during MDS. (A) **1**, (B) **2** and (C) **3**. Numbers in the figure denote times (ns) at which the snap shots were taken.

Analysis of peptide and protein conformation by MDS in DMSO suggests that unfolding appears to depend on the amino acids present. A transmembrane helical peptide that is hydrophobic, does not unfold during 10-ns MDS [28]. Conformational stability can be attributed to solvation of hydrophobic side-chains by DMSO. The beta amyloid peptide Aβ_1-42_ unfolds in the initial 10-ns during MDS in DMSO from a starting structure that is a monomer and in helical conformation [29]. A study on α/β proteins indicate that in DMSO, α-helices unfold first followed by β-sheets [30]. Hydrophobic amino acids would be solvated to a greater extent, that results in conformational stability. NMR data indicate that **1** folds into β-hairpin conformation with well-defined β-turns in DMSO solution with backbone hydrogen bonds [1]. In order to examine whether solvation of hydrophobic residues can result in folding, MDS was carried out in DMSO for **1-3** starting from fully extended structures as in the case of MDS from water.

The data presented in Figure 5 indicate considerable structural fluctuations and no folding into β-hairpin structure, unlike in water.

**Figure 5:**
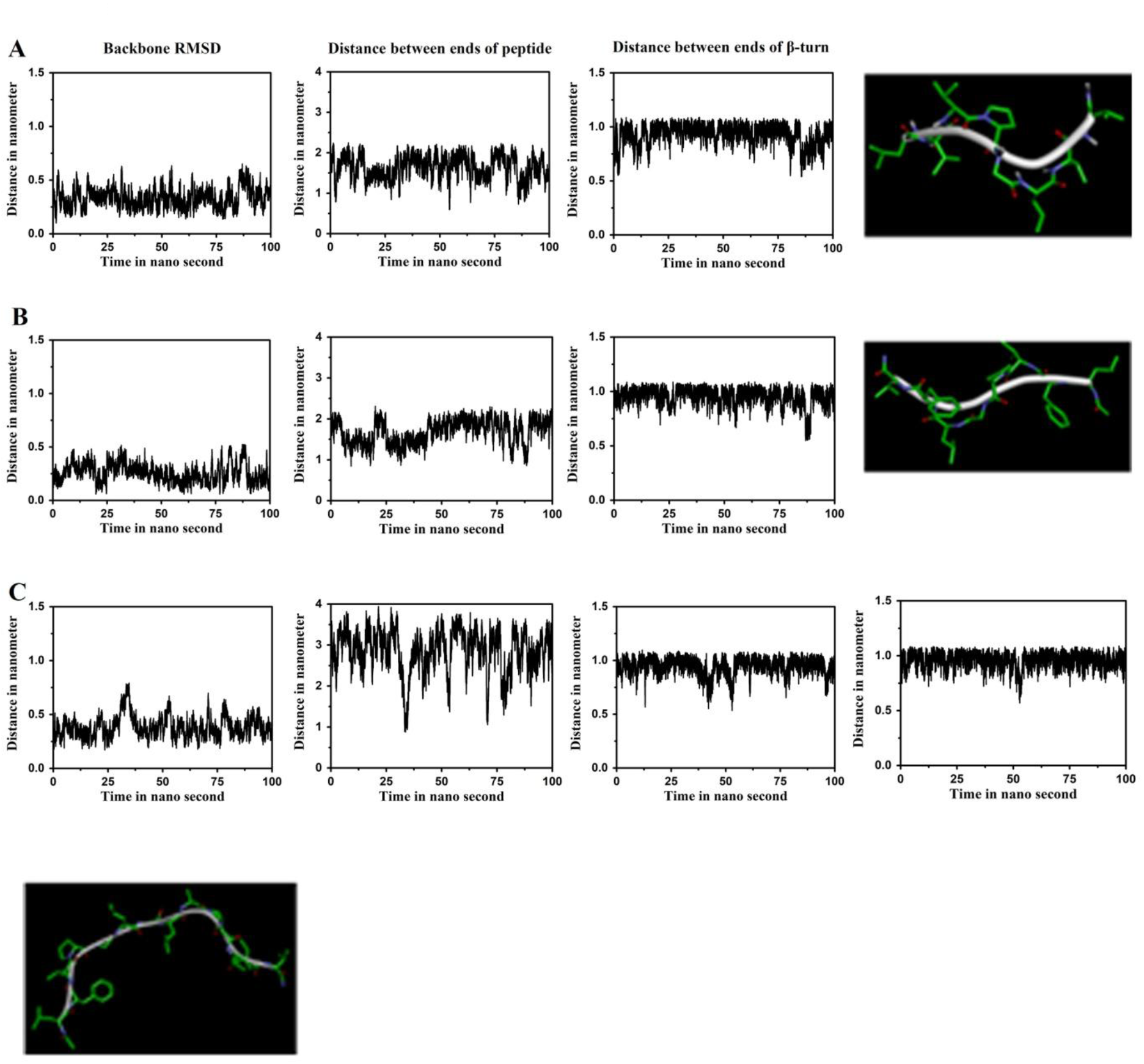
Backbone RMSD, distance between Cα of N and C-terminal amino acids, distance between Cα_i_ and Cα_i+4_ of residues forming β-turns and snapshot of structures at the end of MDS in DMSO for peptides (A) **1**, (B) **2** and (C) **3**.

The basis for the absence of folding was investigated by examining the values of Ф, Ψ during MDS. The data are presented in Figure 6. While the Ф, Ψ values for p are in the region of T2′ β-turn, the G Ф, Ψ values span the entire allowed region for G. This results in the absence of a stable turn and hairpin.

**Figure 6.**
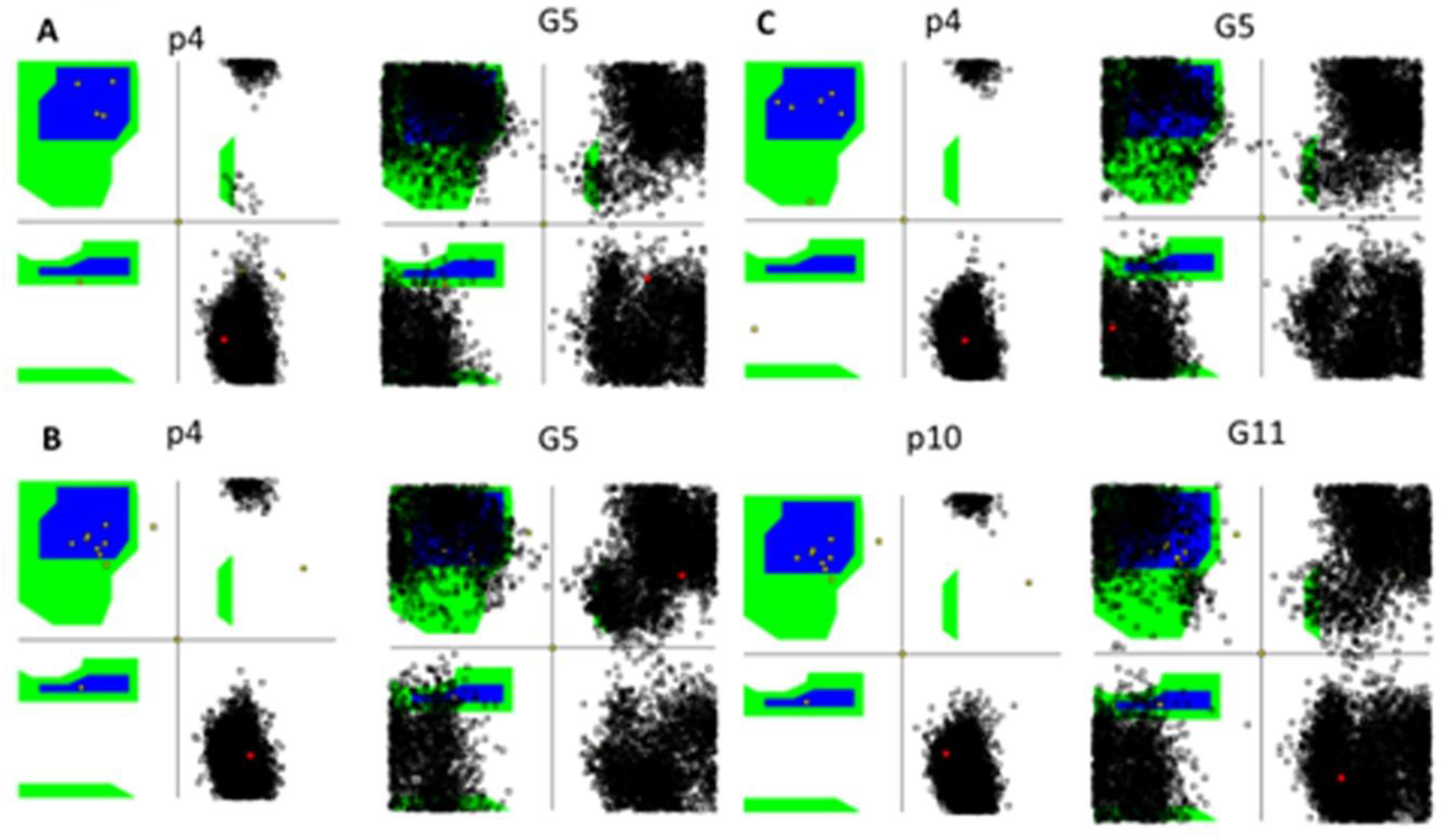
Distribution of Ф, Ψ for p and G in peptides during the course of MDS in DMSO. (A) **1**, (B) **2** and (C) **3**. p4, G5 are D-Pro and Gly at positions 4, 5 in peptides **1-3**. p10, G11 are D-Pro and Gly at positions 10,11 in **3**. Black dots indicate Φ, Ψ values for the indicated amino acids during MDS.

In order to examine the conformational flexibility of i and i+4 residues forming β-turn in these peptides, their Ф, Ψ values were examined (Figure 7). While the sampling of the conformational space in DMSO is more than that in water, it does not sample the entire conformational space available for these amino acids as in the case of Gly.

**Figure 7:**
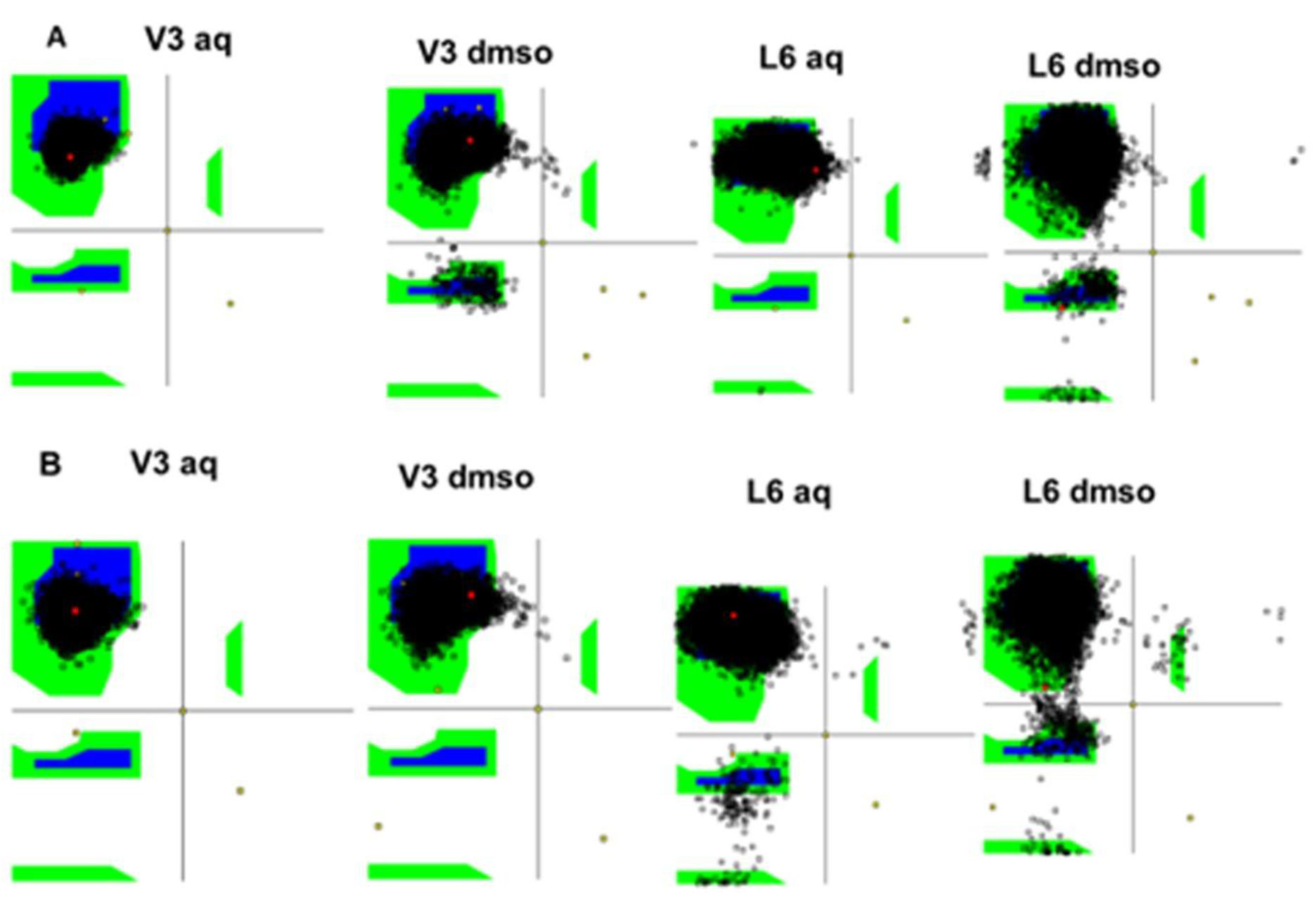
Comparison of Ф, Ψ values of i and i+4 turn residues in peptides during MDS in water (aq) and DMSO. (A) **1** and (B) **2**. V3 and L6 are Val and Leu at positions 3, 6 in peptides **1, 2**. Black dots indicate Φ, Ψ values for the indicated amino acids during MDS.

The effect of DMSO on the conformation of **1** starting from a β-hairpin conformation was examined. The data shown in Figure 8 indicates that unfolding does not occur. Also, the backbone hydrogen bonds are not broken during MDS. Thus, even short, hydrophobic peptides do not unfold in DMSO during MDS similar to a transmembrane helical peptide that is hydrophobic [29].

**Figure 8:**
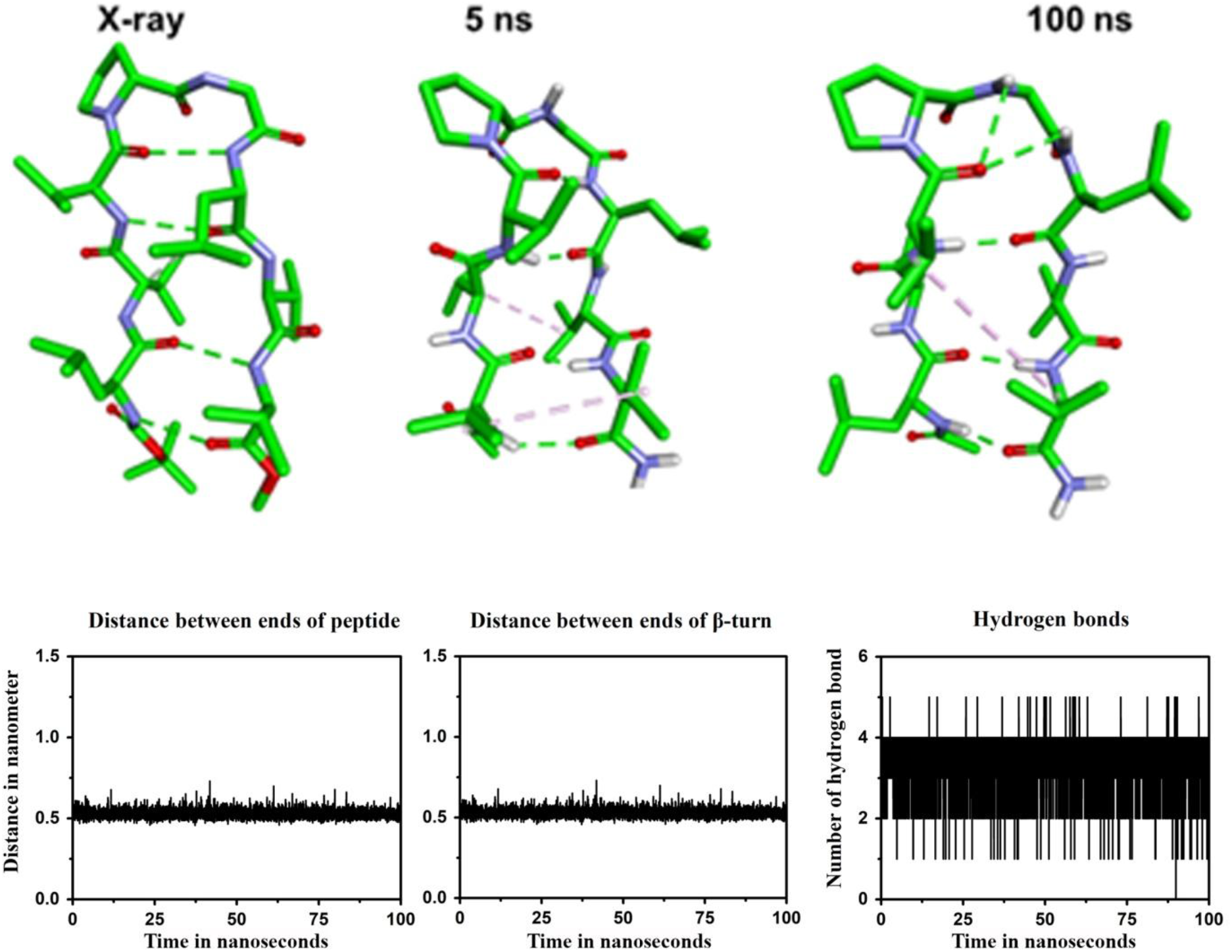
Snapshots (top panels) of **1** starting from folded conformation during MDS in DMSO, distances between N and C-terminal ends, i and i+4 turn residues and hydrogen bonds (bottom panels).

In order to examine whether the absence of folding in pG turn containing peptides in DMSO is due to G, MDS was carried out on peptide **4**, where the turn residues are pA, starting from extended structure. The data shown in Figure 9 indicate folded conformations populated between 25 to 75 ns. However, unfolding occurs after 75 ns. The Ф, Ψ values for p is largely confined to +Ф and –Ψ but Ф, Ψ values for A do not populate the region expected for T2′ β-turns. The T2′ values are populated only in a small fraction of structures. This is also reflected in the snapshots (Figure 10). MDS where the starting structure is in β-hairpin conformation, indicates that unfolding does not occur as in the pG peptide.

**Figure 9:**
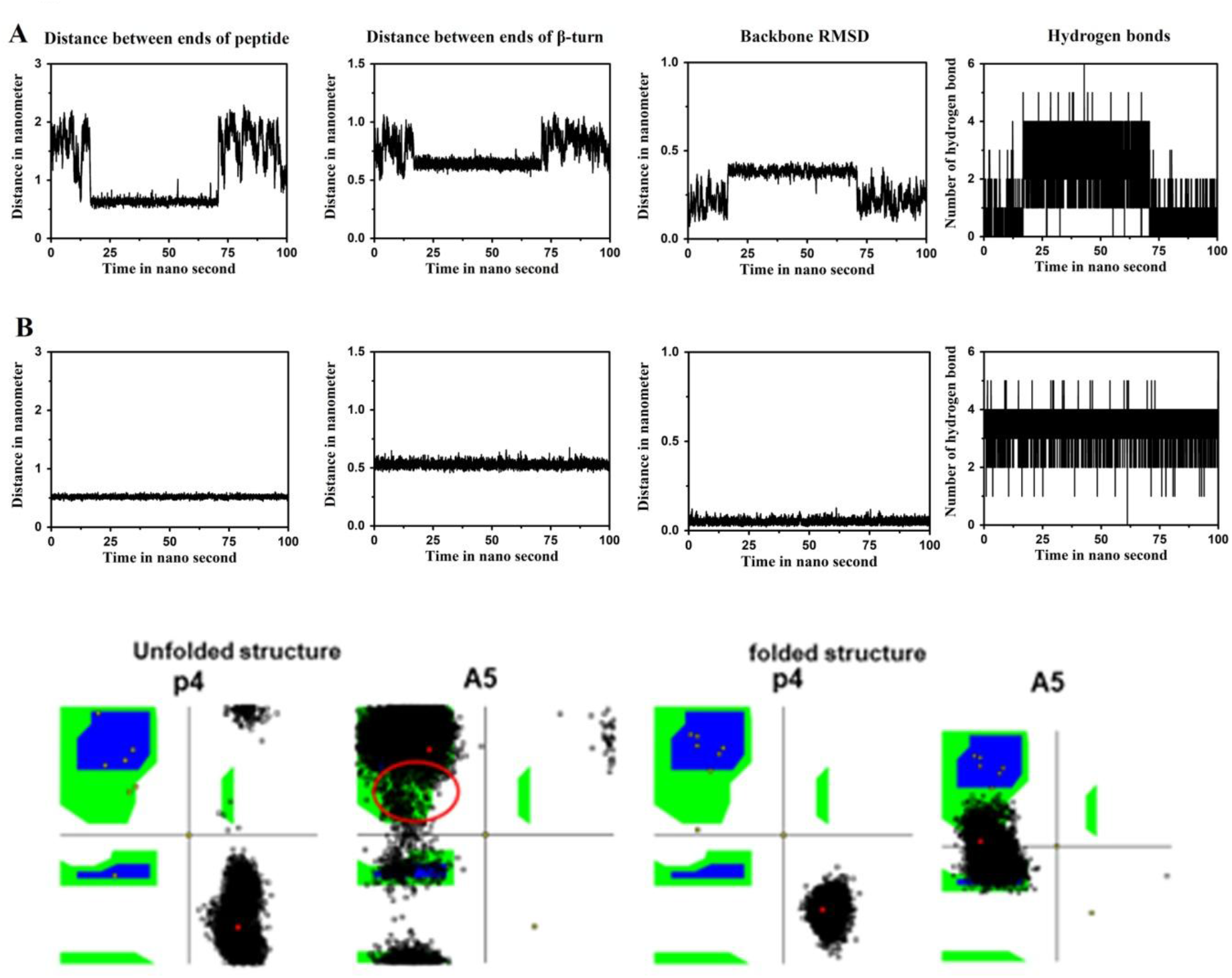
Backbone RMSD, Distance between Cα of N and C-terminal amino acids, distance between Cα_i_ and Cα_i+4_ of residues forming β-turns, backbone RMSD and hydrogen bonds profile in **4** (A) unfolded and (B) folded. Black dots are Ф, Ψ values of p and G during MDS in DMSO where starting structures are unfolded and folded.

**Figure 10:**
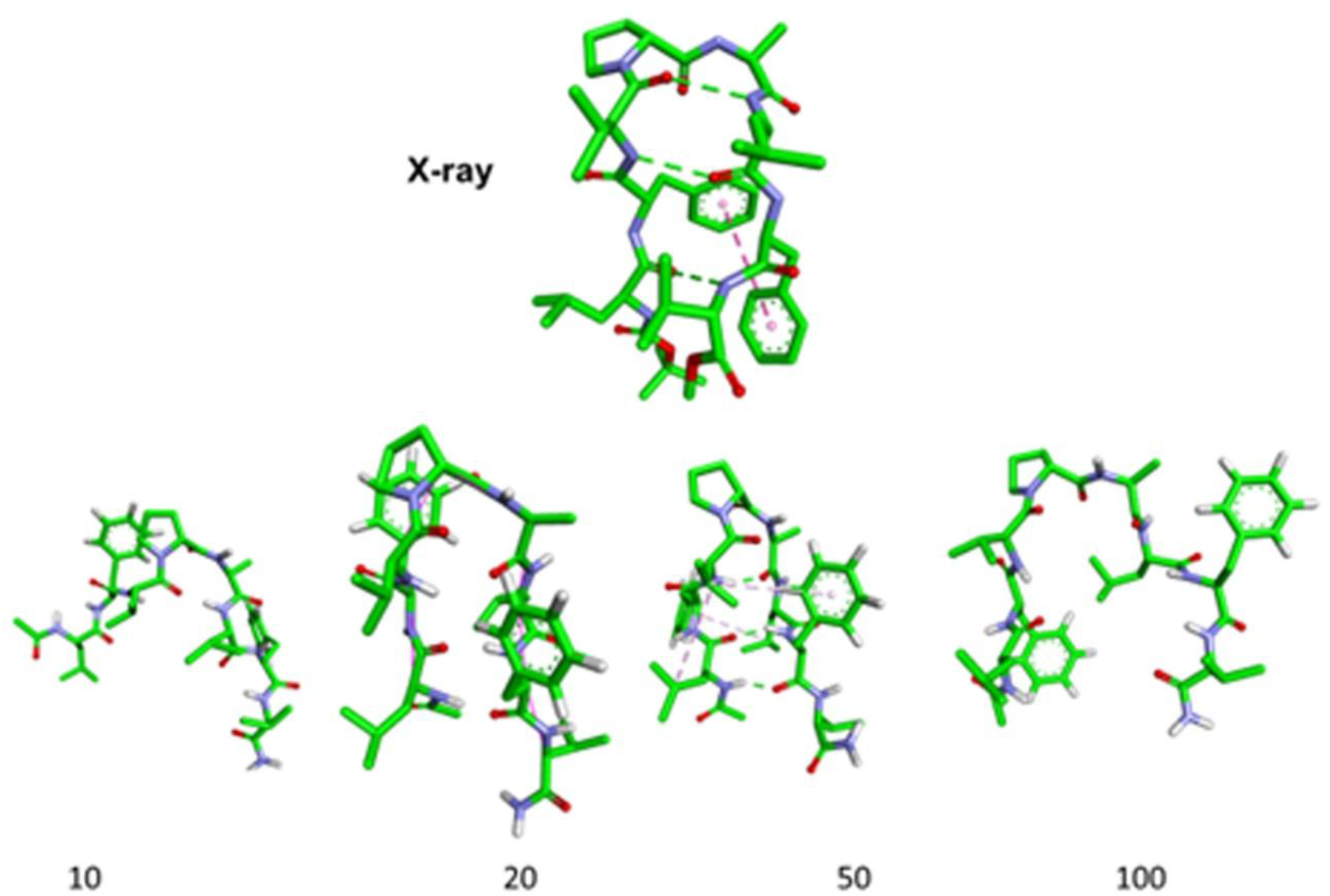
Snapshots of structures for **4** during MDS in DMSO. Numbers indicate times when the snap shots were taken.

NMR studies indicate that several short peptides are unstructured in DMSO (13-16). Detection of a well-defined β-turn and β-hairpin in **1**, is unusual for a linear peptide in DMSO. When the starting structure is in a β-hairpin conformation, unfolding does not occur. Solvation of the hydrophobic side-chains by DMSO results in stabilization of the folded structure. When the starting structure is unfolded, interaction of the hydrophobic side-chains with DMSO and hydrogen bonding of the backbone amide hydrogens with DMSO prevents folding. Water clearly favours folding into β-turn and β-hairpin conformations during MDS, as also observed experimentally for several peptides. Since recent advances in NMR instrumentation would enable studying at uM concentrations, it would be of interest to examine the conformations of **1-4** and related hydrophobic peptides in water.

## Conclusions

Our conformational analysis by MDS indicates that fully protected short peptides composed of only hydrophobic amino acids can fold into β-hairpin conformation in aqueous environment. Experimental studies have indicated that these peptides fold into β-hairpin conformation with a well-defined β-turn in apolar solvents such as CHCl_3_. They have not been investigated by NMR in aqueous medium presumably due to low solubility. NMR studies have suggested that these hydrophobic peptides also fold into β-hairpin conformation in DMSO. When the starting structure is fully extended, the peptides do not fold into β-hairpin conformationin DMSO unlike in aqueous medium, where folding into β-hairpin conformation is observed. However, unfolding is not observed, suggesting that DMSO does not have a denaturing effect on short, hydrophobic peptides.

## Acknowledgements

RN is Indian National Science Academy (INSA) Senior Scientist.

## References

1. Raghothama, S., Awasthi, S., and Balaram, P. β-Hairpin nucleation by Pro-Gly β-turns. Comparison of D-Pro-Gly and L-Pro-Gly sequences in an apolar octapeptide. J. Chem. Soc., Perkin Trans: 1998. 2: 137–144.

2. Das, C., Naganagowda, G.A., Karle, I.L. Balaram, P. Designed β-hairpin peptides with defined tight turn stereochemistry. Biopolymers: 2001. 58: 335–346.

3. Das, C., Raghothama, S. and P. Balaram, P. A designed three stranded β-sheet peptide as a multiple β-hairpin model. J. Am. Chem. Soc. 1998. 120: 5812–5813.

4. Rai, R., Raghothama, S. and Balaram, P. Design of a peptide hairpin containing a central three-residue loop. J. Am. Chem. Soc. 2006. 128: 2675–2681.

5. Tatko, C.D. and Waters, M.L. Selective aromatic interactions in β-hairpin peptides. J. Am. Chem. Soc. 2002. 124: 9372–9373.

6. Chen, P-Y., Lin, C-K., Lee,, C-T. Jan, H. and Chan, S.I. Effects of turn residues in directing the formation of the β-sheet and in the stability of the β-sheet. Protein Sci. 2001. 10: 1794–1800.

7. Searle, M.S., Williams, D.H. and L.C. Packman, L.C. A short linear peptide derived from the N-terminal sequence of ubiquitin folds into a water-stable non-native β-hairpin. Nat Struct Biol. 1995. 2: 999–1006.

8. Maynard, A.J., Sharman, G.J. and Searle, M.S. Origin of β-hairpin stability in solution: structural and thermodynamic analysis of the folding of a model peptide supports hydrophobic stabilization in water. J. Am. Chem. Soc. 1998. 120: 1996–2007.

9. Blanco, F.J., Jimenez, M.A., Herranz, J., Rico, M., Santoro, J. and Nieto,, J.L. NMR evidence of a short linear peptide that folds into a. beta.-hairpin in aqueous solution. J. Am. Chem. Soc. 1993. 115: 5887–5888.

10. Dyson, H.J. and Wright, P.E. Defining solution conformations of small linear peptides. Annu. Rev. Biophys. Biophys. Chem. 1991. 20: 519–538.

11. Karle, I.L., Awasthi, S.K. and Balaram, P. A designed beta-hairpin peptide in crystals. Proc. Natl. Acad. Sci. U S A. 1996. 93: 8189–8193.

12. Teplow, D.B., Preparation of amyloid β-protein for structural and functional studies. Methods Enzymol. 2006. 413: 20–33.

13. Kanyalkar, M., Srivastava, S. and Coutinho, E. Conformation of N-terminal HIV-1 tat (fragment 1–9) peptide by NMR and MD simulations. J Pept Sci. 2001. 7: 579–587.

14. Di Bello, C., Gozzini, L., Tonellato, M,. Grazia Corradini, M., D’Auria, G., Paolillo, L. and Trivellone, E. 500 MHz NMR characterization of synthetic bombesin and related peptides in DMSO-d6 by two-dimensional techniques. FEBS Lett. 1988. 237: 85–90.

15. Chary, K. V., Srivastava, S., Hosur, R. V., Roy, K. B. and Govil, G. Molecular conformation of gonadoliberin using two-dimensional NMR spectroscopy. Eur J Biochem. 1986. 158: 323–332.

16. Pellegrini, M., Gobbo, M., Rocchi, R., Peggion, E., Mammi, S. and D F Mierke, D.F. Threonine6-bradykinin: Conformational study of a flexible peptide in dimethyl sulfoxide by NMR and ensemble calculations. Biopolymers. 1996. 40: 561–569.

17. Karle, I.L., Das, C., and Balaram, P. De novo protein design: crystallographic characterization of a synthetic peptide containing independent helical and hairpin domains. Proc. Natl. Acad. Sci. U S A. 2000. 97: 3034–3037.

18. Aravinda, S., Harini, V.V., Shamala, N., Das, C and Balaram, P. Structure and Assembly of Designed β-Hairpin Peptides in Crystals as Models for β-Sheet Aggregation. Biochemistry 2004, 43: 1832–1846.

19. Vishnyakov, A., Lyubartsev, A.P. and Laaksonen, A. Molecular dynamics simulations of dimethyl sulfoxide and dimethyl sulfoxide− water mixture. J Phys Chem. A, 2001. 105: 1702–1710.

20. James, M., Teemu, A., Roland, M., Szilárd, S., Jeremy, P., Smith, C., Hessa, B and Lindahlad, E. GROMACS: High performance molecular simulations through multi-level parallelism from laptops to supercomputers. SoftwareX, 2015. 1: 19–25.

21. Schmid, N., Eichenberger, A.P., Choutko, A., Riniker, S., Winger, M., Mark, A.E. and van Gunsteren, W. F. Definition and testing of the GROMOS force-field versions 54A7 and 54B7. Eur Biophys J. 2011. 40: 843.

22. Berendsen, H.J.C., Postma, J.P.M., van Gunsteren, W. F. and Hermans, J. Interaction models for water in relation to protein hydration, in Intermolecular forces. 1981, Springer. 331-342. Interaction Models for Water in Relation to Protein Hydration. In: Pullman B. (eds) Intermolecular Forces. The Jerusalem Symposia on Quantum Chemistry and Biochemistry, vol 14. Springer, Dordrecht.

23. Parrinello, M. and Rahman, A. Polymorphic transitions in single crystals: A new molecular dynamics method. J Appl Phys., 1981. 52: 7182–7190.

24. Bussi, G., Donadio, D., and Parrinello, M. Canonical sampling through velocity rescaling. J Chem Phys. 2007. 126: 014101.

25. Darden, T., York, D. and Pedersen, L. Particle mesh Ewald: An N.log (N) method for Ewald sums in large systems. J Chem Phys. 1993. 98: 10089–10092.

26. Miyamoto, S. and Kollman, PA. Settle: An analytical version of the SHAKE and RATTLE algorithm for rigid water models. J Comput Chem. 1992. 13: 952–962.

27. Hess, B., Bekker, H., Berendsen, H.J.C. and Johannes Fraaije, J.G.E.M. LINCS: a linear constraint solver for molecular simulations. J Comput Chem. 1997. 18: 1463–1472.

28. Duarte, A. M. S., van Mierlo, C. P. M. and Marcus A. Hemminga. M. A. Molecular Dynamics Study of the Solvation of an α-Helical Transmembrane Peptide by DMSO. J. Phys. Chem. B 2008, 112: 8664–8671.

29. Lee, M., Chang, H. J., Park, J.Y., Shin, J., Prk, J.W., Choi, J.W., Kim, J.I., Na, S. Conformational changes of A (1–42) monomers in different solvents. Journal of Molecular Graphics and Modelling 2016, 65: 8–14.

30. Roy, S. and Bagchi, B. Comparative Study of Protein Unfolding in Aqueous Urea and Dimethyl Sulfoxide Solutions: Surface Polarity, Solvent Specificity, and Sequence of Secondary Structure Melting. J. Phys. Chem. B 2014, 118: 5691−5697.

